# Nanopore sequencing identifies a higher frequency and expanded spectrum of mitochondrial DNA deletion mutations in human aging

**DOI:** 10.1101/2022.12.05.519134

**Authors:** Amy R. Vandiver, Austin N. Hoang, Allen Herbst, Cathy C. Lee, Judd M. Aiken, Debbie McKenzie, Winston Timp, Jonathan Wanagat

## Abstract

**Background:** Mitochondrial DNA (mtDNA) deletion mutations cause many human diseases and are linked to age-induced mitochondrial dysfunction. Mapping the mutation spectrum and quantifying mtDNA deletion mutation frequency is challenging with next generation sequencing methods. We hypothesized that long-read sequencing of human mtDNA across the lifespan would detect a broader spectrum of mtDNA rearrangements and provide a more accurate measurement of their frequency.

**Results:** We employed nanopore Cas9-targed sequencing (nCATS) to map and quantitate mtDNA deletion mutations and develop analyses that are fit-for-purpose. We analyzed total DNA from vastus lateralis muscle in 15 males ranging from 20 to 81 years of age and substantia nigra from three 20-year-old and three 79-year-old men. We found that mtDNA deletion mutations detected by nCATS increased exponentially with age and mapped to a wider region of the mitochondrial genome than previously reported. Using simulated data, we observed that large deletions are often reported as chimeric alignments. To address this, we developed two algorithms for deletion identification which yield consistent deletion mapping and identify both previously reported and novel mtDNA deletion breakpoints. The identified mtDNA deletion frequency measured by nCATS correlates strongly with chronological age and predicts the deletion frequency as measured by digital PCR approaches. In substantia nigra, we observed a similar frequency of age-related mtDNA deletions to those observed in muscle samples, but noted a distinct spectrum of deletion breakpoints.

**Conclusions:** NCATS-mtDNA sequencing allows identification of mtDNA deletions on a single molecule level, characterizing the strong relationship between mtDNA deletion frequency and chronological aging.

## Introduction

Somatic mitochondrial DNA (mtDNA) deletion mutations are implicated in aging and age-related diseases, but we currently lack a complete picture of the spectrum and quantity of these mutations. Age-induced structural variants of mtDNA, such as deletions, were first reported over 40 years ago. Since that time, mtDNA deletions have been found to contribute to cell dysfunction and cell death [1–3], particularly in post-mitotic tissues, and have been shown to be predictive of age-induced physiological declines [4]. Despite this importance, mapping and quantitation has been hampered by a low frequency in tissue homogenates and a corresponding need for DNA amplification. Numerous methods are available to study mtDNA deletion mutations, but these approaches are often better suited for either mapping or quantitation, rarely both. The inability to quantify and map these mutations limits both their use as potential metrics of biological aging and mechanistic investigation into their origins and consequences.

The multicopy nature and overall structure of mammalian mtDNA has important implications for the study of age-induced structural variants. The human mitochondrial genome is 16,569 base pairs in length compared to ∼6 billion base pairs in the diploid nucleus and is present in 10s to 1000s of copies per diploid nucleus[5,6]. In high copy number tissues such as human skeletal muscle, this means mtDNA makes up about ∼1% of total DNA by mass. To sequence mtDNA with sufficient coverage, studies using homogenate DNA need to account for these factors, most often through amplification or enrichment of mtDNA or removal of nDNA[7,8]. The transposition of mtDNA sequences into nDNA (i.e., termed nuclear mitochondrial DNA segment (NUMTS)) further complicates the study of mtDNA[9]. Structural variants may be of particular importance in the mitochondrial genome because mtDNA lacks introns and has few noncoding regions, so any structural variant is likely to disrupt gene replication, transcription, or translation. Deletion mutations, with sizes ranging from 4 to >15,000 bps, have been the most frequently studied mtDNA structural variant[10,11]. A deletion event may involve flanking direct repeats, but these are not necessary for deletion formation [8,12,13]. Numerous mechanisms are hypothesized to result in deletion formation including defects in both mtDNA replication and double stranded break repair [12].

Diverse methods have been used to study and quantitate somatic mtDNA deletion mutations in aging[14,15]. Southern blot, long-extension PCR, qPCR, and digital PCR approaches have different advantages and disadvantages, which result in differing sensitivities and specificities as well as differing reported mtDNA deletion mutation frequencies[16–19]. In general, mtDNA deletion frequency increases with age in human muscle, but the true mtDNA deletion mutation frequency and spectrum remain unclear because of methodological limitations. Many indirect methods have been used to detect, sequence, and map mtDNA deletion mutations, each of which has inherent biases that limit the spectrum of mutations that can be assessed and complicate quantitation. Single-molecule, direct DNA sequencing offers an opportunity to quantitate and map mtDNA deletion mutations without the need for mtDNA amplification, fragmentation, or enrichment. We have previously demonstrated the utility of nCATS for identifying mtDNA deletions [20], however our prior work focused on a small set of samples at the extremes of lifespan and used a basic analytical approach which did not detect the expected spectrum of larger deletions that have been described in aged mammals.

Here we report our deployment of nanopore Cas9-targeted sequencing (nCATS) to quantify and map mitochondrial DNA deletions from otherwise healthy human skeletal muscle and substantia nigra brain tissue across the lifespan, and provide an initial analytical framework for mtDNA structural variants from long-read sequencing data. We hypothesize that single molecule direct sequencing will yield a higher mutation frequency and a broader spectrum of mutations than previously reported using alternate methods. To address this, we used nCATS in total DNA from human skeletal muscle and substantia nigra and mapped deletions from long-read DNA sequencing data. These data identify deletion mutations at higher frequencies than previously published and involving a broader genomic spectrum. Single molecule DNA sequencing thus provides a novel approach to the quantification of aged-induced mtDNA deletion frequency that does not require DNA amplification, fragmentation, or nDNA removal.

## Results

### Optimization of deletion calling algorithm in simulated data

To identify and map mtDNA deletions in human muscle tissue, we targeted mtDNA with Cas9 targeted sequencing on Oxford Nanopore (mtDNA nCATS) [20]. In mtDNA nCATS, genomic DNA free ends are dephosphorylated and cas9-guided cleavage is then used to introduce new phosphorylated double strand breaks specifically into the mitochondrial genome, exposing novel phosphorylated ends at the sites of interest. Nanopore sequencing adaptors are selectively ligated to the exposed phosphorylated ends, providing selective sequencing of mtDNA.

To determine the efficacy of our initial method for identifying deletions of multiple sizes within the mitochondrial genome, an *in silico* test data set was generated using NanoSim [25]. For this test data, sequencing reads were simulated from template genomes in which deletion events were introduced beginning at bp 5547 and extending 1, 2, 3, 4, 5, 6, 7, 8, 9 or 10 kbp with read lengths and an error profile based on our sequencing data (representative aligned reads in Supplemental Figure 1). When test data was aligned with Minimap2 and deletions called based on the CIGAR sequence of the primary alignment as previously utilized to call mtDNA deletions in nanopore data [20,27], there was a rapid decline in detection with increasing deletion size (Figure 2B). Closer examination of aligned data demonstrated that many reads containing large deletions were aligned as chimeric alignments with a “primary” alignment and a non-overlapping “Supplemental” alignment. To address this, we developed an algorithm to identify deletions between chimeric alignments from Minimap2 (Figure 2A) that are nonoverlapping and contiguous on the query sequence but distant on the reference sequence. Using this approach, we noted increased detection of larger deletions in simulated data (Figure 2B).

To determine if our deletion calling approach was dependent on the choice of aligner, we utilized a parallel method in which chromosome M reads for which query length diverged from aligned length by >500 bp were realigned using BLAST. If this returned two non-overlapping alignments that were contiguous on the query sequence but distant on the reference sequence, a deletion was identified as the space between the two alignments (Figure 2A). Using this method on our simulated data, we observed decreased sensitivity in detecting deletions (Figure 2B), but a similar performance in relationship to deletion size.

### Long-read sequencing of mtDNA identifies mtDNA deletion breakpoints in muscle samples

We applied mtDNA nCATS to 15 human muscle samples obtained from male donors ranging from 26 to 81 years of age (Supplemental Table 1). Prior to sequencing, the frequency of large deletions in each sample was quantified using our validated droplet digital PCR assay [19], which indicated a logarithmically increasing frequency of large deletions with increasing donor age (Supplemental Figure 2). Nanopore sequencing generated an average of 162,947 reads per sample, with a range of 64-85% of reads aligned to the reference mitochondrial genome. Between 82 and 90% of reads aligned to the forward strand of ChrM.

Deletions were first identified through parsing of the CIGAR sequence from the primary alignment of each read as previously reported [20,27]. We identified a total of 10,712 deletion events greater than 100 bp across all 15 samples, ranging from 101 bp to 8131 bp. The previously identified mitochondrial “common deletion,” a deletion event spanning bp 8470-13477, which is associated with Kearns-Sayre syndrome and frequently reported in aged tissue[28], was identified 51 times, representing 5.7% of deletions greater than 2 kb identified (shown in red in Figure 2A). To normalize for differences in read depth in visualizing data, we selected a random subsample of 30,000 reads from each sample for plotting (Figure 2A). Next, to understand the frequency of deletions per molecule of mtDNA, we calculated the frequency of deletions of specific minimum sizes per mtDNA read. To determine what size of deletions most significantly correlates with chronological age, we calculated the correlation coefficient for the frequency of deletions greater than each size threshold versus chronological age (Figure 2B).

The highest correlation to chronologic age was observed for deletions greater than 3 kbp (R2=0.83, Figure 2C). The abundance of deletions greater than 3 kbp per read detected using nCATS was correlated with the log-transformed deletion frequency as obtained by droplet digital PCR (R^2^=0.38, Figure 1E).

**Figure 1:**
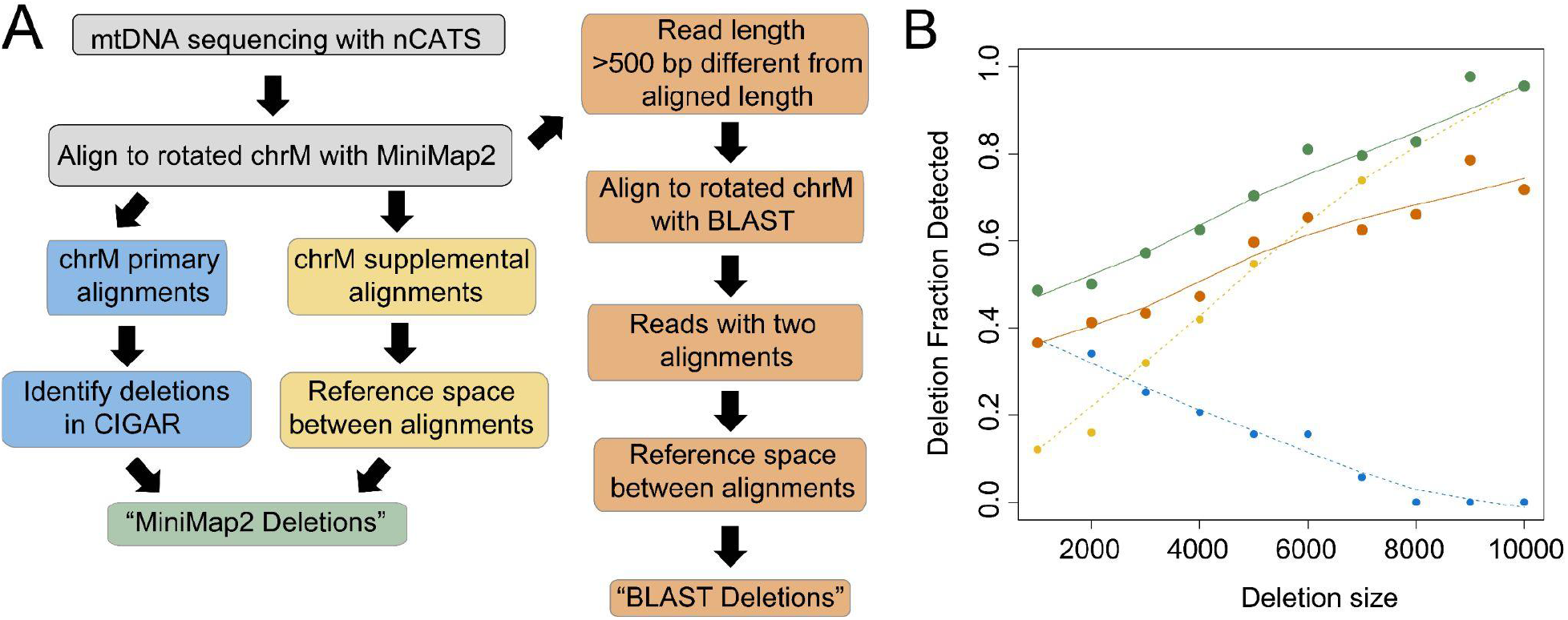
Optimization of mtDNA deletion calling algorithms using *in silico* test data. A) Schematic of algorithms used for identifying large mtDNA deletions. B) Fraction of expected deletions identified versus size of expected deletion in simulated data set. Results of primary alignment algorithm shown in blue, supplemental alignment algorithm in yellow, total MiniMap2 alignment in green, BLAST alignment algorithm in orange.

Given the increased ability to identify larger deletions when considering chimeric alignments, we next applied the algorithm optimized on simulated data to identify deletions using Minimap2 chimeric alignments and combined these with the deletions identified in primary alignments. With this combined set we identified a total of 12,621 deletion events with sizes ranging from 138 bp to 14,146 bp (deletions greater than 2 kbp in a subset of 30k reads per sample shown in Figure 3A). In the total set, the “common deletion” was detected 100 times, representing 4.18% of deletions greater than 2 kbp. The strongest correlation (R^2^=0.84) of deletions per read to chronological age was observed when considering deletions >2 kbp(Figure 3B-C). This averages to a frequency of 4.3 × 10^−5^ for 25-year-old individuals and increases 59 fold to 2.5 × 10^−3^ for 75-year-old individuals. The frequency of deletions per mtDNA read correlates strongly with ddPCR deletion frequency (R^2^=0.66), with a higher deletion frequency noted in sequencing data than ddPCR at all ages (Figure 3E). Comparable results were obtained when we applied the BLAST deletion calling algorithm. Using this approach, we detected a total of 10,810 deletions events ranging from 101 bp to 14045 bp across all samples (deletions greater than 2 kbp in a subset of 30k reads per sample shown in Figure S3A). A strong correlation (R^2^=0.66) between deletion frequency and age (Figure S3 B-C) and deletion frequency and ddPCR deletion frequency was also observed using this approach (Figure S3 D-E). In addition to changes in deletion frequency, the mean size of deletions was observed to increase significantly with age (p<0.01) (distribution shown in Figure 4A).

**Figure 2:**
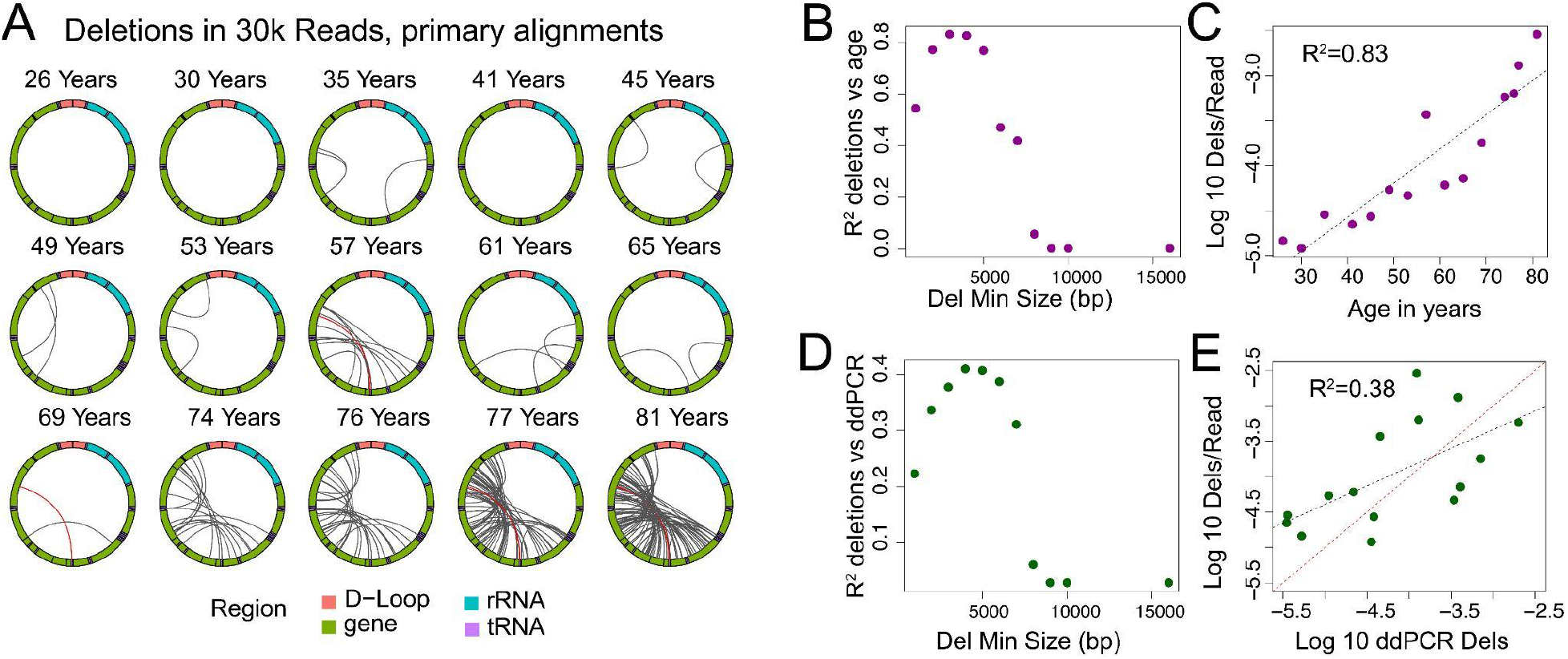
Large mtDNA deletions in primary alignments. A) Localization of deletions >2000 bp identified in CIGAR sequence of primary alignments in a random subset of 30,000 reads per sample. The mitochondrial genome is depicted in a clockwise orientation with lines linking the start and end of each identified deletion. The human mitochondrial “common” deletion is highlight in red where identified. B) R^2^ correlation coefficient for log transformed number of identified deletions per mitochondrial read versus age plotted versus the minimum deletion size cutoff used in calculating correlation. C) Log transformed frequency of deletions >3 kbp per mitochondrial read versus age. D) R^2^ correlation coefficient for log transformed number of identified deletions per mitochondrial read versus log transformed deletion frequency by ddPCR plotted versus the minimum deletion size cutoff used in calculating correlation. E) Log transformed frequency of deletions >3kbp per mitochondrial read versus log transformed ddPCR deletion frequency. The red dashed line is the line of identity.

**Figure 3:**
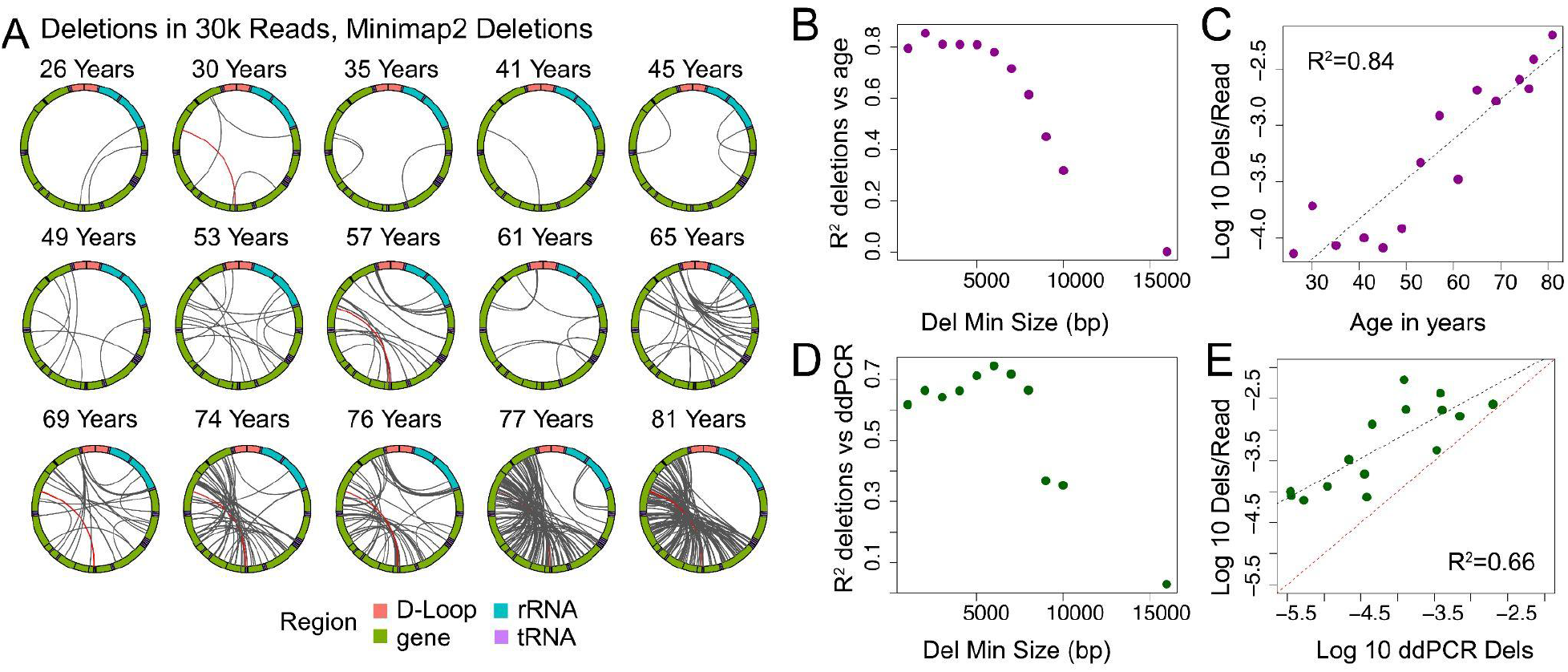
Large mtDNA deletions mapped using combined Minimap2 alignments. A) Localization of deletions >2000 bp identified using the Minimap2 algorithm in a random subset of 30,000 reads per sample. The mitochondrial genome is depicted in a clockwise orientation with lines linking the start and end of each identified deletion. The human mitochondrial “common” deletion is highlight in red where it was detected. B) R^2^ correlation coefficient for log transformed number of identified deletions per mitochondrial read versus age plotted versus the minimum deletion size cutoff used in calculating correlation. C) Log transformed frequency of deletions >2 kbp per mitochondrial read versus age. D) R^2^ correlation coefficient for log transformed number of identified deletions per mitochondrial read versus log transformed deletion frequency by ddPCR plotted versus the minimum deletion size cutoff used in calculating correlation. E) Log transformed frequency of deletions >2kbp per mitochondrial read versus log transformed ddPCR deletion frequency. Red dashed line is the line of identity.

**Figure 4:**
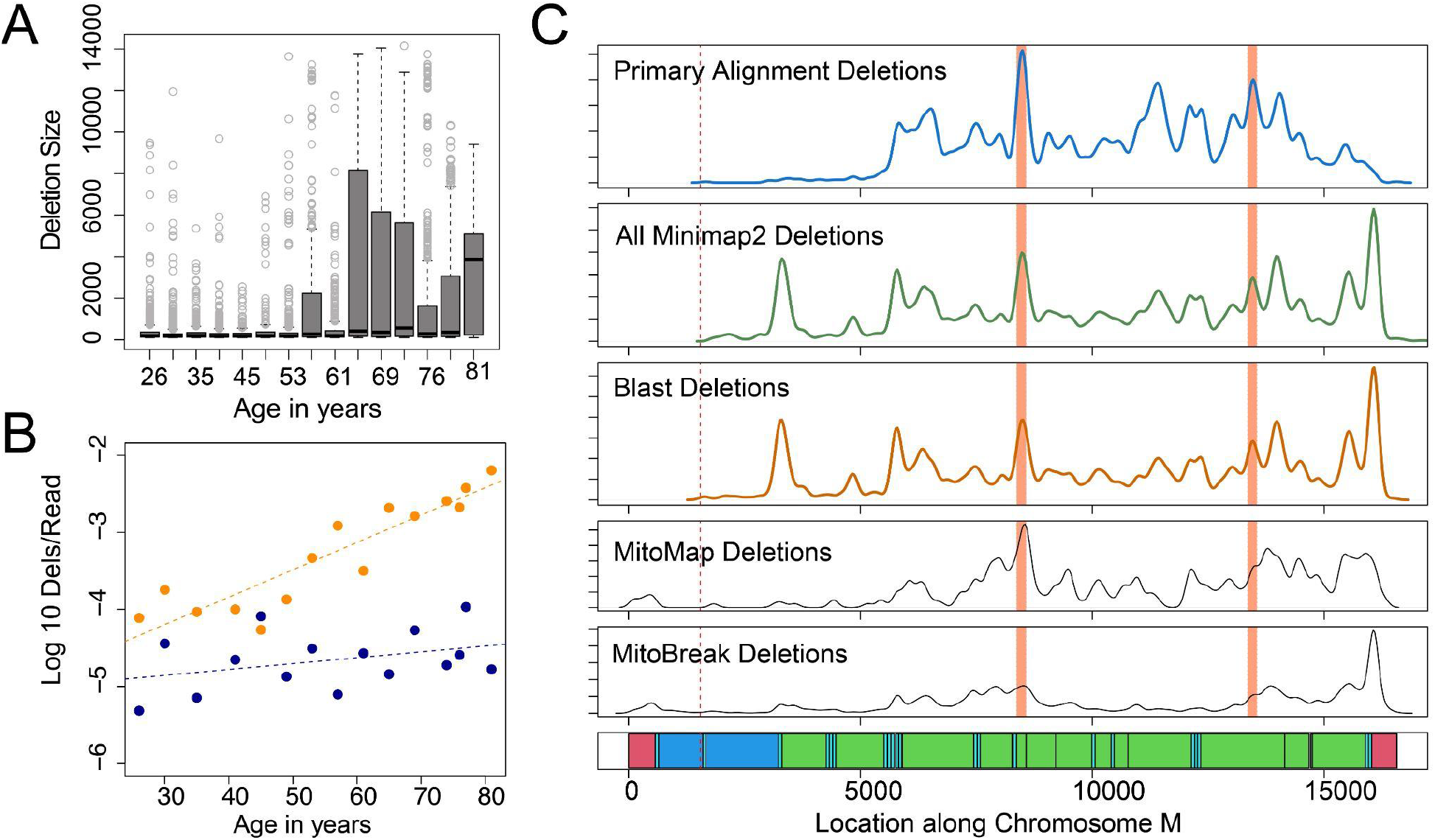
Distribution of mtDNA deletions across the mitochondrial genome. A) Boxplots depicting distribution of mtDNA deletion size versus age of sample. B) Log transformed frequency of deletions >2 kbp per mitochondrial read versus age, plotted separately for deletions contained entirely within the minor arc (dark blue) and deletions involving the major arc (orange). C) Density of deletion breakpoints identified using primary alignment algorithm (blue), combined Minimap2 deletions (green) and blast alignment deletions (dark orange) as compared to all deletion breakpoints reported in MitoMap and MitoBreak databases. Bottom panel shows annotation of the mitochondrial genome. D-loop regions shown in red, rRNA shown in dark blue, tRNA shown in light blue, coding regions shown in green. Cas9 cut site shown in dotted lines. Location of human common deletion breakpoints shown in light red rectangles.

### Long-read sequencing maps mtDNA deletions across mitochondrial genome

The ability to directly map deletion breakpoints in a less biased manner with nCATS mtDNA sequencing provides new opportunities to understand the distribution of mtDNA deletion breakpoints across the mitochondrial genome. To begin understanding these data, we compared the distribution of deletion breakpoints identified using our methods to deletion breakpoints identified through other methods in the Mitomap[29] and MitoBreak[10] databases (Figure 4C). Utilizing permutation testing, deletion breakpoints identified in primary alignments were not significantly different from deletion breakpoints identified in both Mitomap and Mitobreak (p=0.70, p=0.24). When Minimap2 chimeric alignments were also considered, the breakpoint distribution was significantly distinct from deletion breakpoints identified in Mitomap (p=0.03) but not from those identified in MitoBreak (p=0.52). Notably, no significant difference was identified between deletion breakpoints identified using Minimap2 and Blast alignments (p=0.93, Figure 4C). Intriguingly, while deletions within the “minor arc” spanning bp 408-5746, were noted, the frequency of deletions identified entirely within the minor arc did not change significantly with age (p=0.2), while the frequency of deletions involving the major arc correlates strongly with age (p<0.001). This pattern was observed considering deletions greater than 2 kbps (Figure 4B) and deletions of all sizes (Supplemental Figure 5).

### Large mtDNA deletions increase with age in human substantia nigra

To assess the applicability of our algorithm to a non-muscle tissue with known age-related mtDNA changes, we next used nCATS to obtain long-read sequencing of mtDNA from human substantia nigra. For this analysis, we sequenced DNA isolated from substantia nigra from three 20-year-old male donors and three 79-year-old male donors. Nanopore sequencing generated a mean of 123,784 reads per sample, with a range of 9% to 75% of reads aligned to the reference mitochondrial genome. In these samples, we determined the location of mtDNA deletions using Minimap2 alignments. Using this analysis, we identified a total of 1945 deletion events, ranging from 101 to 13,909 bp (Deletions >2 kbp identified in a subset of 12k reads per sample shown in 5A). We noted a significantly increased number of deletions >2 kbp per read in the 79-year-old as compared to 20-year-old donors (Welch’s t-test, p=0.038). We next examined the distribution of deletion breakpoints identified in substantia nigra samples as compared to muscle samples and the published databases. A significant difference was noted between the distribution of deletion breakpoints identified in muscle and substantia nigra samples (P<0.01) with a decrease in deletion breakpoints identified at tRNA-T (bp position 15,888-15,953) in substantia nigra tissue. Despite this difference in distribution, the frequency of mtDNA deletions per mtDNA read was similar between substantia nigra and muscle samples at each age time point (Figure 5C). All deletion breakpoint locations are provided in Supplemental File 1.

**Figure 5:**
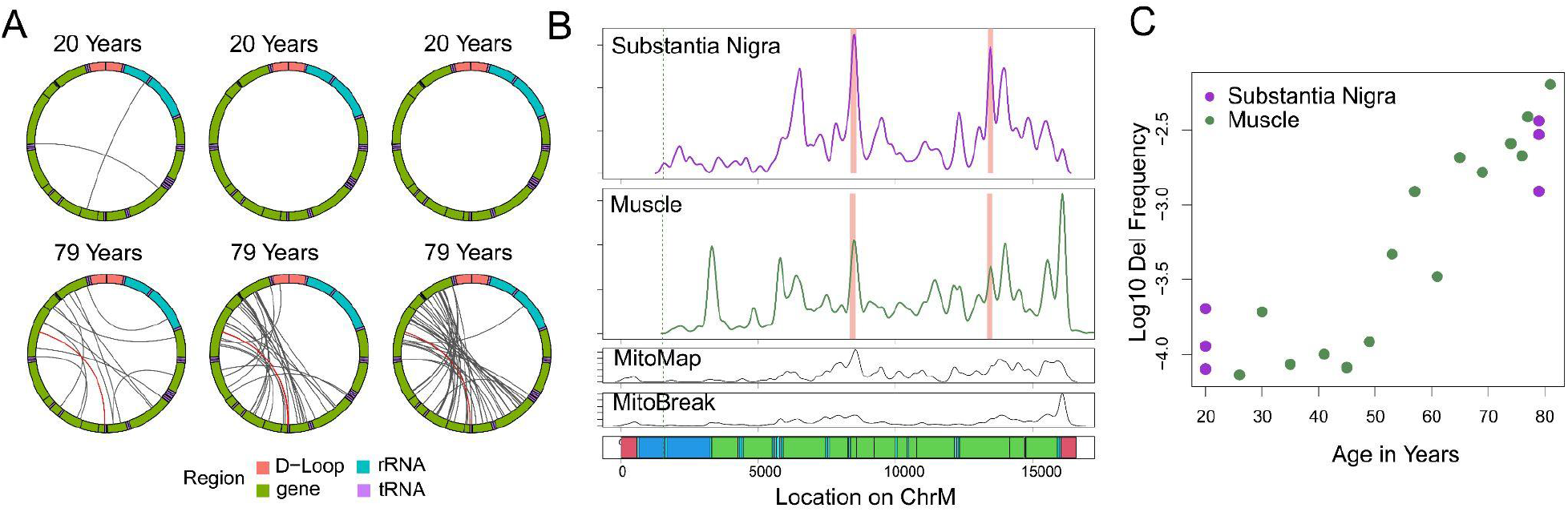
Mapping of large mtDNA deletions in human substantia nigra. A) Localization of deletions >2000 bp identified using minimap2 alignments in a random subset of 12,000 reads per sample. The mitochondrial genome is depicted in a clockwise orientation with lines linking the start and end of each identified deletion. The human mitochondrial “common” deletion is highlighted in red where identified. B) Density of deletion breakpoints identified in human substantia nigra (top panel), as compared to those identified in human muscle tissue (second panel), the MitoMap database (third panel) and the MitoBreak database (fourth panel). Bottom panel shows the annotation of the mitochondrial genome. D-loop regions shown in red, rRNA shown in dark blue, tRNA shown in light blue, coding regions shown in green. Cas9 cut site shown in dotted lines. Location of human common deletion breakpoints shown in light red rectangles. C) Log 10 transformed frequency of mtDNA deletions >2000 bp in total chromosome M reads vs age for muscle (green) and substantia nigra (purple) samples.

Next, to test the utility of our method on a tissue with low expected mtDNA deletion frequency, we utilized nCATs to obtain long read mtDNA sequencing from two human placenta samples. Nanopore sequencing generated an average of 79644 reads per sample, with 24.4% and 38.7% of reads aligned to the reference mitochondrial genome. Using Minimap2 alignments, only 7 deletions greater than 2 kbp were detected across both samples, consistent with the expected low level of age-related mtDNA change (Supplemental Figure 4).

## Discussion

In multiple tissues across the human lifespan, we deploy nCATS (nanopore Cas9-targeted sequencing) to provide an amplification-free mapping and quantitation of age-induced mtDNA deletion frequency. Using this approach and a novel mapping algorithm to address chimeric alignments, we find mtDNA deletion mutation frequencies that are higher than generally reported for aging human skeletal muscle, an expanded scope of deletion breakpoints and a tissue specific spectrum of deletions. We observe that the mapping algorithms used have important effects on the spectrum of deletions called and their quantitation. Finally, we demonstrate an age associated shift in the deletion events detected: the mean length of deletions detected increases with age of sample and the proportion of deletions detected within the minor arc decreases.

These findings expand our current understanding of age-induced mtDNA deletion mutations. Many different approaches have been used to detect and quantitate mtDNA deletion mutations including Southern blot, PCR, qPCR, long-extension PCR, digital PCR, and NGS[8,16–19]. Because of the excess of nuclear DNA in total DNA samples, most approaches to mtDNA deletion mutation detection leverage some type of enrichment, including mitochondrial isolation[30], nuclease digestion of nDNA[8,31], and preferential amplification of mtDNA[7], each of which comes with biases. The nCATS approach avoids DNA amplification, decreasing some of these biases and allowing new insights. In our work, we find up to ∼10-fold higher mutation frequencies using the nCATS approach and a strong, exponential correlation between mutation frequency and age, demonstrating the relevance of the identified changes. These data support our hypothesis that the nCATS approach detects an expanded population of deletion events.

Detecting a larger population of deletion events also reveals a broader spectrum of mtDNA deletion mutation breakpoints. In previous studies, breakpoints were observed predominantly within the mitochondrial major arc spanning from base pair 5747 to 407 [32], although breakpoints spanning both major and minor arcs [33] and entirely within the minor arc have also been reported [8]. One major arc deletion, known as the mitochondrial “common deletion”, has been frequently reported in tissue homogenate studies, but has not been detected in single muscle fiber or single cell studies. In our work, we observe the presence of the common deletion as well as many of the previously reported deletion breakpoints, validating the utility of our method for breakpoint detection. However, in addition to observation of these previously reported events, we observe novel deletion breakpoints, including those within the mitochondrial minor arc. This finding leads to two intriguing observations: first, we note that while minor arc deletions are present, their frequency does not increase with age as seen with major arc deletions, suggesting possible different mechanisms for development and amplification of minor versus major arc deletions. Second, we note a distinct distribution of deletion breakpoints in substantia nigra versus muscle tissue samples, indicating possible tissue specificity in the mechanism of deletion formation or amplification. While these findings have potential impacts for the study of mtDNA deletions, we note that a full understanding of the intricacies of deletion distribution with age and between tissues will require larger sample sizes across the lifespan.

This work demonstrates the potential of nCATS-mtDNA for mapping and quantitating age-induced deletion mutations, laying a foundation to deepen the understanding of these mutations in human aging and relevant clinical settings. We note that the current work is limited in scope, and thus does not fully elucidate the broad potential for studying human aging. Specifically, this initial study was limited to male samples and only 15 skeletal muscle and 6 substantia nigra samples. We have only male subjects at this point due to our focus on physical function with age in older US veterans. MtDNA deletion mutation frequency, on average, is lower in women than in men[4] using dPCR methodology. We hypothesize that nanopore sequencing will reveal similar sex differences in skeletal muscle deletion frequency across the human lifespan. A benefit of our current sample set is the orthogonal ddPCR data from the same samples, which add rigor to our findings of age-induced increases in mtDNA deletion mutation frequency in human muscle. The second main limitation is the lack of established analysis algorithms for mtDNA deletion detection in long-read sequencing. The algorithms presented here represent first efforts in aligning and mapping mtDNA structural variants. We expect that emerging data on long-read mtDNA sequencing will accelerate the development and validation of improved algorithms for these analyses, akin to the continued evolution of pipelines to detect such variants in NGS data [34,35]. The third main limitation is the use of a single cut site located in the minor arc for mtDNA cleavage. The selection of this single site was based on our current understanding of the known distribution of age- and disease-induced mtDNA deletion mutations[10,33]. The data from this study demonstrate that a broader spectrum of mtDNA deletions exists and thus, using different cut sites may further alter the deletion breakpoint distribution in future studies. Optimization of the analysis algorithms would be prudent before adding additional cut sites or attempts at sample multiplexing.

To further the utility of mtDNA nCATS for use in human studies, future work will focus on expansion and diversification of the data set and continued optimization of analytical methods. The data generated will continue to serve as a template for optimization of analytical pipelines to streamline mtDNA deletion mutation mapping. Applying mtDNA nCATS to more human tissues and including samples from both sexes will extend the relevance and generalizability of the data set. Optimization of the experimental and analytical pipelines would also afford opportunities for more formal validation of performance characteristics for nCATS including limit of detection, precision, and specificity[36]. Here we demonstrate that long-read, direct sequencing of human mtDNA enhances the detection and quantitation of age-induced mtDNA deletion mutations. These data indicate that there is more to learn regarding the frequency and spectrum of these mutations in aging tissues.

## Conclusions

NCATS-mtDNA sequencing allows identification of mtDNA deletions on a single molecule level, characterizing the strong relationship between mtDNA deletion frequency and chronological aging in multiple tissues. These data offer new insight into the frequency and distribution of age-associated mtDNA deletions, providing a foundation for further study into the mechanism of mtDNA deletion formation and accumulation and the use of mtDNA deletions as a metric of tissue aging.

## Materials and methods

### Human subjects

De-identified muscle biopsy specimens were collected as part of a VA Merit Award, “Testosterone, inflammation and metabolic risk in older Veterans” and NIH R01DK090406 (PI: Cathy C. Lee, MD). Use of the human specimens for this study was approved by the UCLA Institutional Review Board (Protocol #18-001547) and the University of Alberta Health Research Ethics Board #00084515. The biopsy samples were obtained from the vastus lateralis muscle of 15 male subjects ranging in age from 20 to 81 years [21]. Personnel analyzing human muscle biopsy samples were blinded to subject age. Subject ages were only revealed after analyses were completed. Substantia nigra samples were obtained from the NIH NeuroBiobank, all subjects were men with no known neurologic disease.

### DNA isolation and quality control

Tissue samples were powdered under liquid nitrogen using a mortar and pestle. Approximately 25 mg of powdered muscle was used for DNA isolation, performed by proteinase K and RNase A digestion, phenol/chloroform extraction, and ethanol precipitation, as previously described [22]. Quality control of the DNA was performed via Nanodrop (Nanodrop 2000, Thermo Scientific; Waltham, MA), Qubit (2.0, Invitrogen; Carlsbad, CA), and gel electrophoresis. Quality control cutoffs for total DNA included A260/280 > 1.8; A260/230 > 1.8.

### Cas9 cleavage, library preparation, and nanopore sequencing

A custom guide RNA sequence targeting human mtDNA: ACCCCTACGCATTTATATAG was used in accordance with previously described methods[20]. Pre-complexed Alt-R CRISPR-Cas9 single guide RNA (IDT) was diluted to a concentration of 10 μM and combined with HiFi Cas9 Nuclease V3 (IDT, cat 1081060) in CutSmart Buffer (NEB, cat B7204). Cas9 cleavage and library preparation was performed in accordance with previously described methods[23]. Approximately 3 μg of input DNA was dephosphorylated with alkaline phosphatase (ONT, cat SQK-CS9109) in CutSmart Buffer (NEB, cat B7204). Following enzyme inactivation of alkaline phosphatase, 10 μL of 333 nM Cas9-gRNA complex was combined with the dephosphorylated DNA, dATP (ONT, cat SQK-CS9109), and Taq DNA polymerase (ONT, cat SQK-CS9109). After Cas9 cleavage and dA-tailing, sequencing adaptors were ligated to target DNA ends and library purification was performed using the Oxford Nanopore Technologies Cas9 Sequencing Kit (ONT, cat SQK-CS9109). Samples were run on a MinION flow cell (v9.4.1) using the MinION Mk1C sequencer. Sequencing and basecalling was operated using the MinKNOW software (v21.10.8).

### Sequence and structural rearrangement analysis

Analysis was done using python version 3.6.15 and R version 4.0.2.

#### Reference genome

The chromosome M reference from HG38 was rotated by removing the base pairs prior to our cut site (1:1547) and concatenating these to the end of the reference sequence in order to account for the expected start and end of our sequenced DNA and prevent aberrant deletion calling

#### Base calling

Base calling was performed using Guppy (v4.5.4, release 4/20/21) on MinKnow (v4.2.8). Reads meeting a QC threshold of 5 were included in analysis.

#### Alignment

All sequences passing quality control were aligned to the rotated mitochondrial genome using Minimap2[24], version 2.24.

#### Simulated data

Test data was generated using NanoSim[25] (version 3.0.2). A representative run of our muscle data, was used as a reference for read profile generation to identify read lengths and quality parameters on which simulated data was based. 500 reads were simulated from 10 different template genomes. Each genome contained a deletion beginning at bp 4000 of the rotated genome, extending either 1, 2, 3, 4, 5, 6, 7, 8, 9 or 10 kbp. After read generation, all 5000 reads were combined for subsequent alignment and deletion calling in order to generate a simulated data set with and even distribution of deletions of each size.

#### Deletions in primary alignment

Deletions were identified from primary alignments by parsing the CiGAR sequence of each ChrM primary alignment using GenomicAlignments package in R (version 1.26.0) [26] to identify deletions greater than 100 bp. Deletions within 300 bp were combined to a single deletion event.

#### Deletions in supplemental alignments

Alignments to ChrM with the 2048 flag were extracted. Positional information and span across the reference genome for each supplemental alignment and the paired primary alignment was determined using RSamtools (version 2.6.0). For reads <17000 bp in which the primary and supplemental alignments were each >200 bp, were contiguous within 300 bp on the query sequence but distant on the reference sequence, and primary and supplemental alignments overlap by less than 50 bp on query and reference sequences,deletions were identified as the position between the primary and supplemental alignments on the reference genome.

#### Deletion calling using BLAST

For reads with primary alignment to ChrM, the total read length was compared to the length of the primary alignment on the reference genome using RSamtools. For reads in which the read length was >500 bp shorter or longer than the aligned length, the sequence was realigned to the rotated reference genome using BLAST (Basic Local Alignment Search Tool) aligner via rBLAST (version 0.99.2). For reads <17000 bp with two non-overlapping alignments in BLAST that were each >200 bp, were contiguous within 300 bp on the query sequence but distant on the reference sequence, deletions were identified as the position between the alignments.

#### Density analysis

Distribution of deletion breakpoints across the mitochondrial genome were compared using a permutation test of equality through the sm package (version 2.2-5.7) with a smoothing parameter of 50.

Analysis code is available at https://github.com/amyruthvandiver/Nanopore_Tissues

## Declarations

### Ethics approval and consent to participate

De-identified muscle biopsy specimens were collected as part of a VA Merit Award, “Testosterone, inflammation and metabolic risk in older Veterans” and NIH R01DK090406 (PI: Cathy Lee, MD). Use of the human specimens for this study was approved by the UCLA Institutional Review Board (Protocol #18-001547) and the University of Alberta Health Research Ethics Board #00084515.

### Consent for publication

Not applicable

### Availability of data and material

The data that support the findings of this study are available from the corresponding author upon reasonable request and through SRA, under Bioproject PRJNA844096. Analytical code is available at https://github.com/amyruthvandiver/Nanopore_Tissues.

### Authors’ contributions

Study designed and conceived by AV, ANH, and JW. Sample preparation was performed by ANH. Data analyses were performed by AV. All authors contributed to writing and revising the manuscript.

## Acknowledgements

Not applicable

## Supplementary materials

Supplemental Figure 1: Integrative genome viewer view of simulated data set.

Supplemental Figure 2: Droplet digital PCR deletion frequency increases exponentially with age.

Supplemental Figure 3: Identification of mtDNA deletions in skeletal muscle samples using Blast algorithm.

Supplemental Figure 4: Frequency of minor arc and major arc deletions versus age for all deletions.

Supplemental Figure 5: Identification of mtDNA deletions in placenta tissue. Supplemental Table 1: Annotation of tissue samples used in analysis.

Supplemental File 1: Deletion breakpoints identified in each tissue.

