## Supplemental Figure 1, Supplemental Figure 2, Supplemental Figure 3, Supplemental Figure 4, Supplemental Figure 5 for "Nanopore sequencing identifies a higher frequency and expanded spectrum of mitochondrial DNA deletion mutations in human aging"

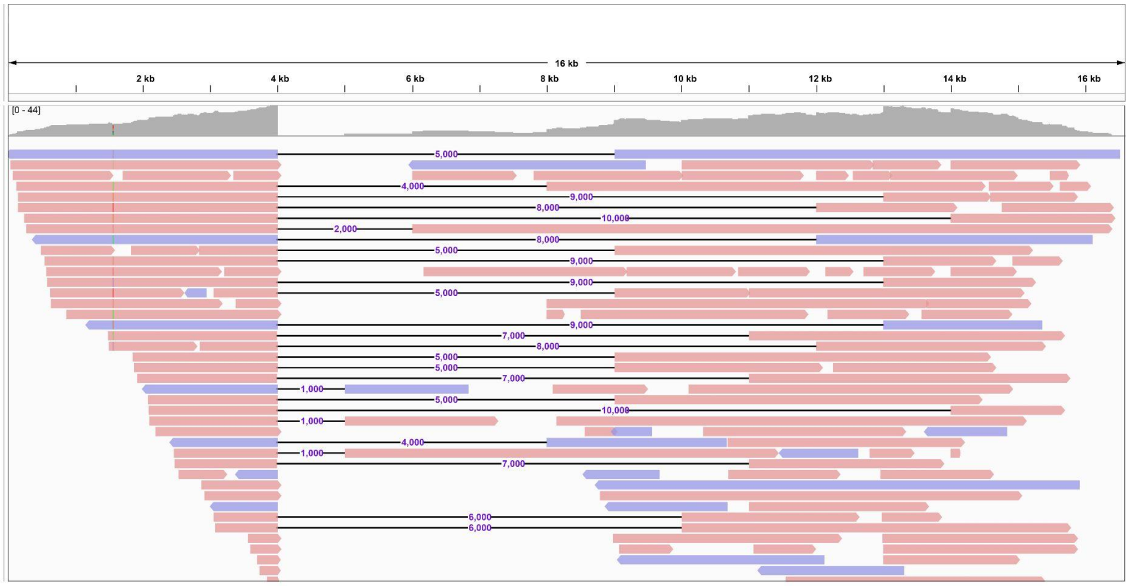


**Supplemental Figure 1: Integrated genome viewer of simulated data set.** Top panel depicts mitochondrial genome coordinates. Middle panel depicts coverage depth across the mitochondrial genome. Bottom panel depicts individual simulated reads as pink (forward strand) or purple (reverse strand) bars.

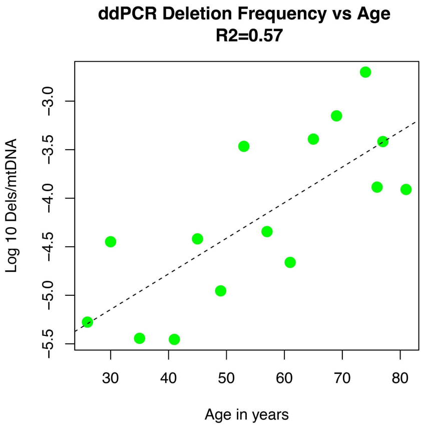


**Supplemental Figure 2: Droplet digital PCR deletion frequency increases exponentially with age.** Shown are the log 10 transformed droplet digital PCR measured mtDNA deletion frequencies (y-axis) versus sample age.

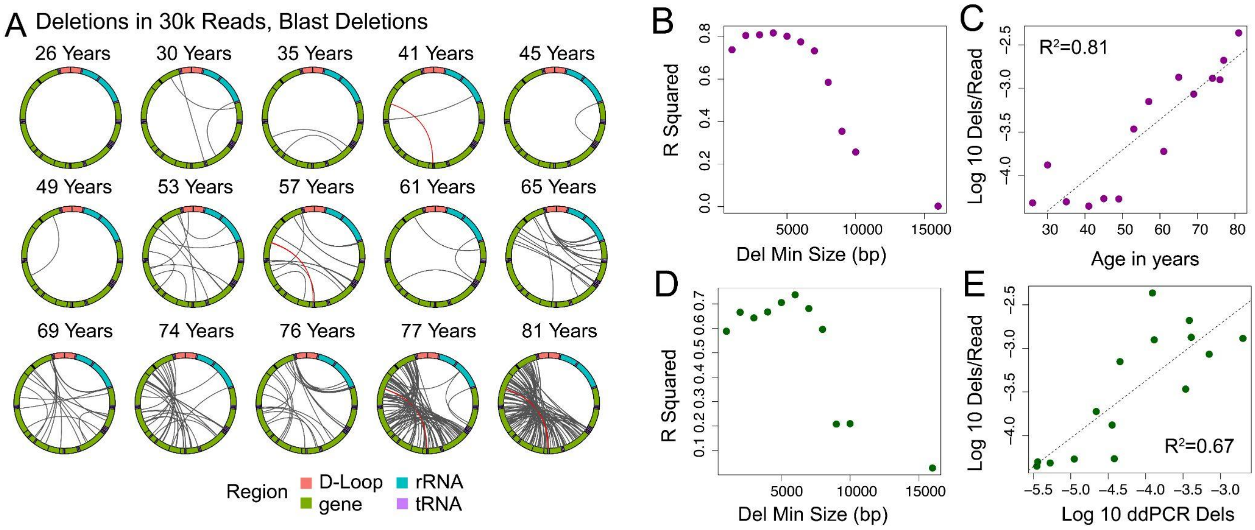


**Supplemental Figure 3: Identification of mtDNA deletions in skeletal muscle samples using Blast algorithm.** A) Localization of deletions >2000 bp identified as chimeric alignments using the Blast aligner in a random subset of 30,000 reads per sample. The mitochondrial genome is depicted in a clockwise orientation with lines linking the start and end of each identified deletion. The human mitochondrial "common" deletion is highlight in red where identified. B) $R^2$ correlation coefficient for log transformed number of identified deletions per mitochondrial read versus age plotted versus the minimum deletion size cutoff used in calculating correlation. C) Log transformed frequency of deletions >2 kbp per mitochondrial read versus age. D) $R^2$ correlation coefficient for log transformed number of identified deletions per mitochondrial read versus log transformed deletion frequency by ddPCR plotted versus the minimum deletion size cutoff used in calculating correlation. E) Log transformed frequency of deletions >2kbp per mitochondrial read versus log transformed ddPCR deletion frequency. Red indicates line of identity.

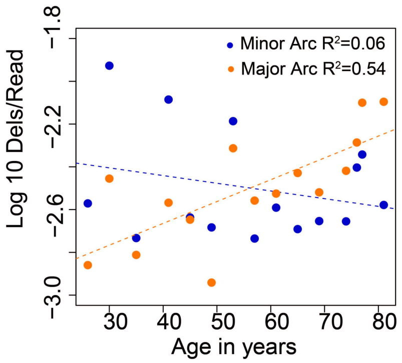


**Supplemental Figure 4: Frequency of minor arc and major arc deletions versus age for all deletions.** Log transformed frequency of all identified deletions per mitochondrial read versus age, plotted separately for deletions contained entirely within the minor arc (dark blue) and deletions involving the major arc (orange).

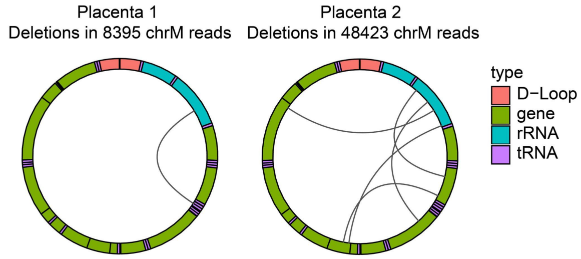


**Supplemental Figure 5: Large mtDNA deletions in placenta samples.** Localization of deletions >2000 bp identified in placenta samples. The mitochondrial genome is depicted in a clockwise orientation with lines linking the start and end of each identified deletion.
